# Engineered Trimeric ACE2 Binds and Locks “Three-up” Spike Protein to Potently Inhibit SARS-CoVs and Mutants

**DOI:** 10.1101/2020.08.31.274704

**Authors:** Liang Guo, Wenwen Bi, Xinling Wang, Wei Xu, Renhong Yan, Yuanyuan Zhang, Kai Zhao, Yaning Li, Mingfeng Zhang, Xingyue Bao, Xia Cai, Yutang Li, Di Qu, Shibo Jiang, Youhua Xie, Qiang Zhou, Lu Lu, Bobo Dang

**Affiliations:** Center for Infectious Disease Research, Zhejiang Provincial Laboratory of Life Sciences and Biomedicine, Key Laboratory of Structural Biology of Zhejiang Province, School of Life Sciences, Westlake University, Hangzhou, Zhejiang, China; Institute of Biology, Westlake Institute for Advanced Study, Hangzhou, Zhejiang Province, China; Key Laboratory of Medical Molecular Virology (MOE/NHC/CAMS), School of Basic Medical Sciences and Biosafety Level 3 Laboratory, Fudan University, Shanghai 200032, China

## Abstract

SARS-CoV-2 enters cells via ACE-2, which binds the spike protein with moderate affinity. Despite a constant background mutational rate, the virus must retain binding with ACE2 for infectivity, providing a conserved constraint for SARS-CoV-2 inhibitors. To prevent mutational escape of SARS-CoV-2 and to prepare for future related coronavirus outbreaks, we engineered a *de novo* trimeric ACE2 (T-ACE2) protein scaffold that binds the trimeric spike protein with extremely high affinity (K_D_ < 1 pM), while retaining ACE2 native sequence. T-ACE2 potently inhibits all tested pseudotyped viruses including SARS-CoV-2, SARS-CoV, eight naturally occurring SARS-CoV-2 mutants, two SARSr-CoVs as well as authentic SARS-CoV-2. The cryo-EM structure reveals that T-ACE2 can induce the transit of spike protein to “three-up” RBD conformation upon binding. T-ACE2 thus represents a promising class of broadly neutralizing proteins against SARS-CoVs and mutants.

---

Coronavirus disease 2019 (COVID-19) caused by SARS-CoV-2 has resulted in a severe global pandemic. Following SARS-CoV, SARS-CoV-2 is yet another emergent beta-coronavirus threatening human health ^1^. SARS-CoV-2 and SARS-CoV (SARS-CoVs) are very similar by sharing 79.5% sequence identity ^1^, similar spike protein structures ^2–4^ and same cell surface receptor angiotensin converting enzyme II (ACE2) ^1,5^. However, seventeen years after the severe acute respiratory syndrome (SARS) pandemic, no targeted vaccines or therapeutics have been approved for SARS, some of which might have held promise for treating COVID-19. Many neutralizing antibodies against SARS-CoV-2 are currently being urgently developed ^6–10^, and some of these might become available later this year or next year. However, RNA viruses are known to have higher mutation rates ^11,12^. Many SARS-CoV-2 mutations have already been identified such as D614G ^13–16^. This means that resultant mutation strains could make current SARS-CoV-2 neutralizing antibodies ineffective in the future. The appearance of COVID-19 after SARS indicates the likely emergence of other coronavirus pandemics in the future. Thus, therapeutics broadly effective against SARS-CoV-2 and mutants, even other SARS-CoV-2-related coronaviruses, are highly desirable. Both SARS-CoV-2 and SARS-CoV bind ACE2 for cell entry, suggesting that SARS-CoV-2 mutants and future related coronaviruses are also likely to bind ACE2. Therefore, proteins engineered based on wild-type ACE2 could serve as the most broadly neutralizing proteins against these viruses and will be least likely to face mutational escape.

The biological function of ACE2 further supports using ACE2 decoy proteins to treat patients infected with SARS-CoVs. Coronavirus infection, or even spike protein binding, can cause shedding of ACE2 from cell surface resulting in a decreased level of ACE2 expression and accumulation of plasma angiotensin II ^17–19^. This phenomenon is closely related to acute lung injury ^17,20–22^. Replenishing soluble ACE2 could alleviate acute respiratory distress syndrome (ARDS) ^17,21–23^. In fact, it has been shown that recombinant soluble ACE2 could inhibit infection from SARS-CoVs in cell assays and organoids ^24–26^. One clinical trial (NCT04335136) was also registered to use recombinant ACE2 to treat COVID-19. However, recombinant soluble ACE2 inhibits viral infection at relatively high concentrations ^24,26–28^, therefore, it may not be an optimal inhibitor. Engineered ACE2 bearing multiple mutations and ACE2-Ig have better spike protein binding affinities and better virus inhibition activities ^25,27–29^. Since spike proteins of SARS-CoVs function as trimers ^2–4^, we reasoned that an engineered trimeric ACE2 protein could potentially bind up to three receptor binding domains (RBD) on the spike protein, which would dramatically increase binding affinity through avidity and thus potently inhibit SARS-CoVs (fig. S1).

To develop such trimeric ACE2 proteins, we chose a C-terminal domain of T4 fibritin (foldon) ^30,31^, or a three helix bundle (3HB) ^32,33^, as trimerization motifs since these have been successfully demonstrated to form stable protein trimers ^3,30,31^. We then looked at the reported SARS-CoVs spike protein structures to determine the most appropriate linker between trimerization motifs and ACE2 ^3,34–38^. It has been found that SARS-CoV spike protein mostly adopts one- or two-up RBD conformations and can engage one or two ACE2 monomers ^34,35^. On the other hand, a very small population of SARS-CoV spike protein can have three-up RBD conformation to bind three ACE2 monomers. SARS-CoV-2 structures mostly have closed conformation or one RBD in the up position ^3,36,37^. From these structural analyses, we estimated that distances between RBDs on the same spike protein could range from 60 Å to 100 Å when they are in the up positions. Moreover, structures from SARS-CoV viral particle revealed there are about 100 spike protein trimers displayed on the 100 nm diameter viral particle surface giving interspike protein distance around 200 Å ^4,39,40^. To retain the possibility for intraspike or interspike avidity, we chose a flexible (GGGGS)5 linker, or a more rigid (EAAAK)5 linker, to construct trimeric ACE2 ^41^. We used wild-type ACE2 peptidase domain (1-615) to construct all trimeric ACE2 decoy proteins. Linkers were inserted after ACE2, followed by the trimerization motifs. We therefore constructed four trimeric ACE2 proteins: ACE2-flexible-3HB, ACE2-rigid-3HB, ACE2-flexible-foldon and ACE2-rigid-foldon. In addition, we constructed two trimeric ACE2 proteins with a short linker GGGS (ACE2-short-3HB, ACE2-short-foldon) and a monomeric ACE2 as control proteins (fig. S2).

## Results

We first used ELISA assay to determine binding affinities between ACE2 proteins and the prefusion stabilized trimeric SARS-CoV-2 spike protein ectodomain (S-ECD) (**Fig. 1**) ^3^. ACE2 monomer binds S-ECD with IC_50_ of 27 nM. For trimeric ACE2 proteins, we saw significant binding affinity enhancement. We found that the rigid linker constructs had the highest binding affinities, including ACE2-rigid-3HB and ACE2-rigid-foldon, which both bound S-ECD with IC_50_ of 30 pM. Trimeric ACE2 proteins with short linkers had lower binding affinities (fig. S3).

**Fig. 1.**
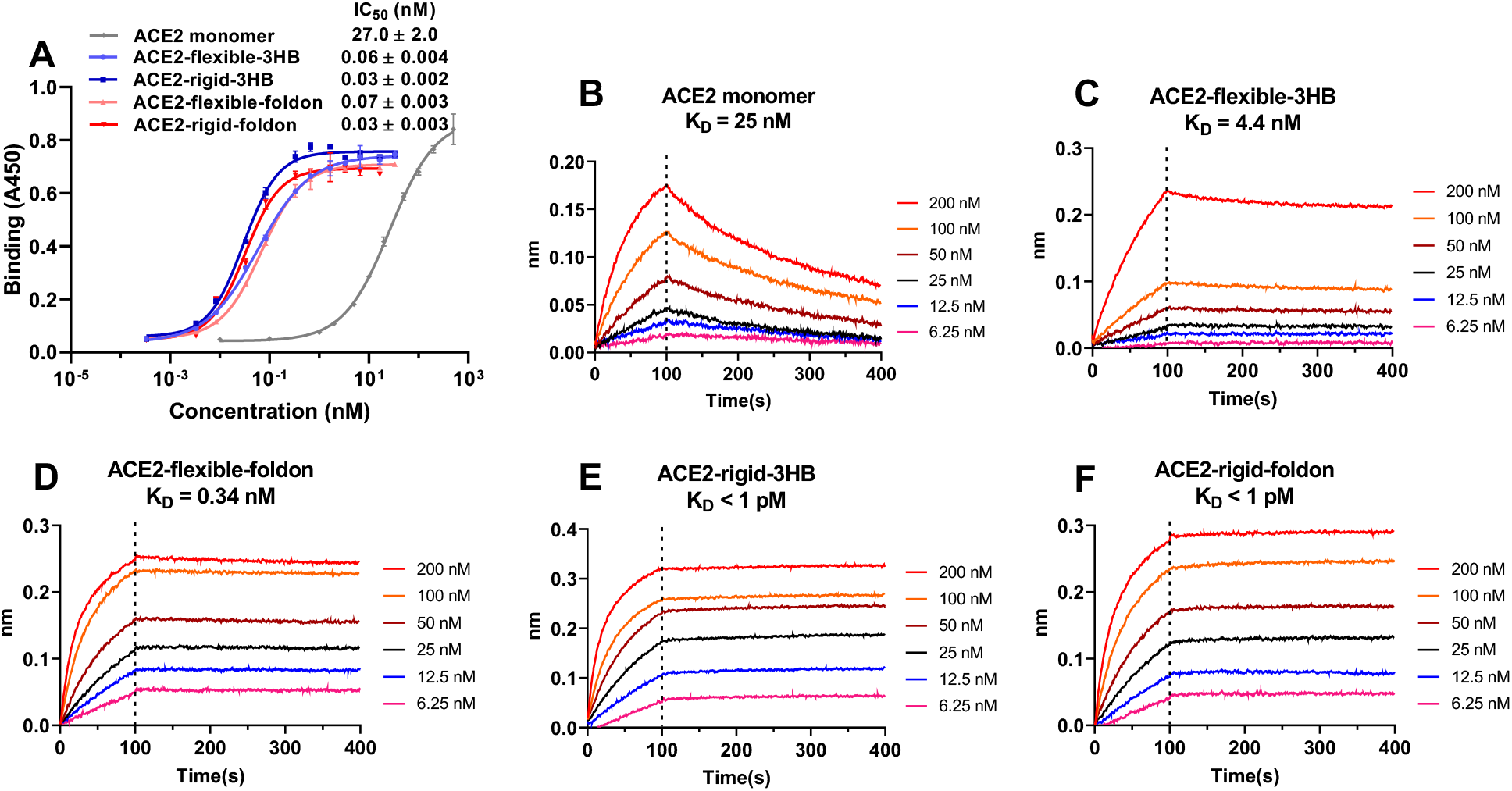
Binding affinities measurements between ACE2 proteins and SARS-CoV-2 spike protein ectodomain (S-ECD). **(A)** Binding affinities measured using ELISA assay. **(B-F)** Binding affinities measured using biolayer interferometry.

We further analyzed ACE2 proteins binding using biolayer interferometry (ForteBio Octet RED96) (**Fig. 1**). S-ECD was biotinylated with NHS-PEG8-Biotin and was loaded on streptavidin coated sensors at about 25% saturation to avoid artificial intermolecular avidity, followed by titration of engineered ACE2 decoy proteins as analytes. K_D_ for ACE2 monomer/S-ECD was 25 nM, agreeing very well with previously published results ^3^. For trimeric ACE2 proteins, we again observed dramatically increased binding affinities. K_D_ for ACE2-flexible-3HB/S-ECD was 4.4 nM while K_D_ for ACE2-flexible-foldon/S-ECD went down to 0.34 nM. Both ACE2-rigid-3HB and ACE2-rigid-foldon bound S-ECD extremely tight, with K_D_ < 1 pM. Further decreasing loading of S-ECD on streptavidin sensors did not affect ACE2 proteins binding, suggesting intramolecular avidity binding between trimeric ACE2s and S-ECD (fig. S4). The massive binding affinity enhancement for ACE2-rigid-3HB and ACE2-rigid-foldon also indicates that spike protein probably has at least two RBDs in the up position upon binding.

Next, we assessed the inhibitory activities of these trimeric ACE2 decoy proteins using SARS-CoV-2 and SARS-CoV pseudotyped viruses in an infection assay of Huh-7 cells (**Fig. 2**). ACE2 monomer can only inhibit SARS-CoV-2 pseudotyped virus at high concentration with IC_50_ > 50 nM. As expected, trimeric ACE2 with flexible linkers showed much better inhibitory activities. ACE2-flexible-3HB inhibited SARS-CoV-2 infection with IC_50_ of 3.46 nM, while ACE2-flexible-foldon had better inhibitory activity with IC_50_ of 1.58 nM. Rigid-linker trimeric ACE2 proteins again displayed the highest inhibitory activities. ACE2-rigid-3HB and ACE2-rigid-foldon showed similar IC_50_’s of 0.40 nM and 0.48 nM, respectively. Short-linker trimeric ACE2 proteins showed no dramatically improved inhibitory activities compared with ACE2 monomer, even though they have higher binding affinities than ACE2 monomer (fig. S5). Similar results were observed with SARS-CoV pseudotyped virus inhibition. (**Fig. 2**). ACE2 monomer had weak inhibitory activity with IC_50_ > 50 nM, while ACE2-rigid-foldon had the best inhibitory activity with IC_50_ of 2.41 nM. Thus, the ACE2-rigid-foldon construct is the most potent trimeric ACE2, which is designated T-ACE2 hereafter.

**Fig. 2.**
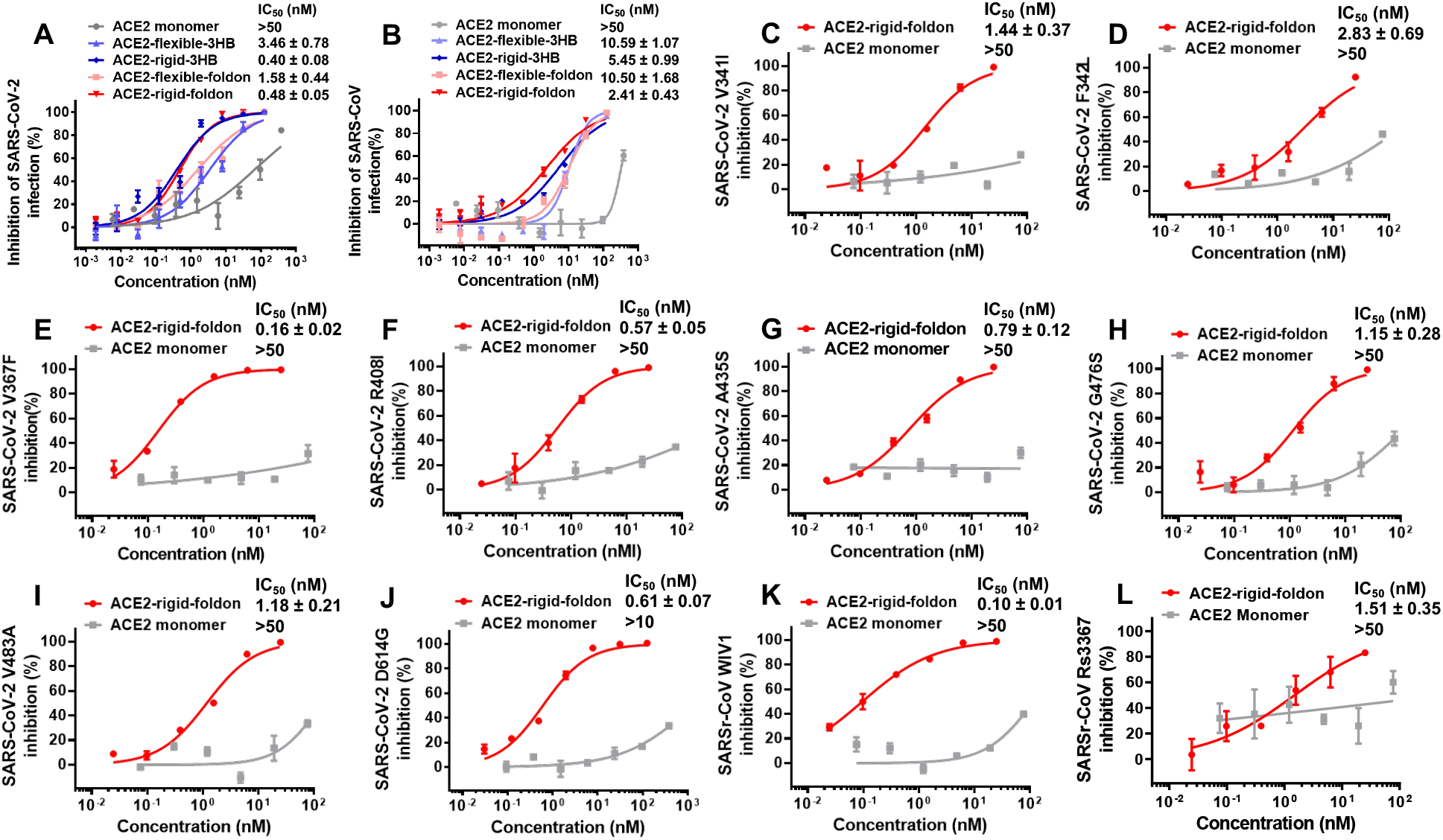
ACE2 proteins inhibition of SARS-CoVs pseudotyped viruses (n=3). **(A)** ACE2 proteins inhibition of SARS-CoV-2. **(B)** ACE2 proteins inhibition of SARS-CoV. **(C-J)** ACE2-rigid-foldon (T-ACE2) inhibition of SARS-CoV-2 mutants. **(K-L)** ACE2-rigid-foldon (T-ACE2) inhibition of SARSr-CoVs WIV1 and Rs3367.

We then asked whether T-ACE2 could also inhibit SARS-CoV-2 mutants and related coronaviruses (**Fig. 2**). We tested T-ACE2 inhibitory activities on eight naturally occurring SARS-CoV-2 mutants, including seven RBD domain mutations ^14,16^, D614G mutation ^13^ and two SARSr-CoVs (WIV1 and Rs3367). We found that T-ACE2 could potently inhibit all these viruses at low nM to sub-nM IC_50_ concentrations (**Fig. 2**). Future identified novel mutations and related coronaviruses are unlikely to escape ACE2. The observed inhibitory activities prompt us to speculate T-ACE2 will have a high probability to inhibit many of these novel mutants if not all.

We further tested T-ACE2 inhibition of authentic SARS-CoV-2 virus (**Fig. 3**). Vero E6 cells were infected with authentic SARS-CoV-2, and inhibitory efficacy was then evaluated using quantitative real-time (qPCR) and confirmed with visualization of virus nucleoprotein (N protein) through immunofluorescence microscopy. Importantly, we found that T-ACE2 could also potently inhibit authentic SARS-CoV-2 with IC_50_ of 1.88 nM, which agreed well with our binding affinity and pseudotyped virus inhibition results.

**Fig. 3.**
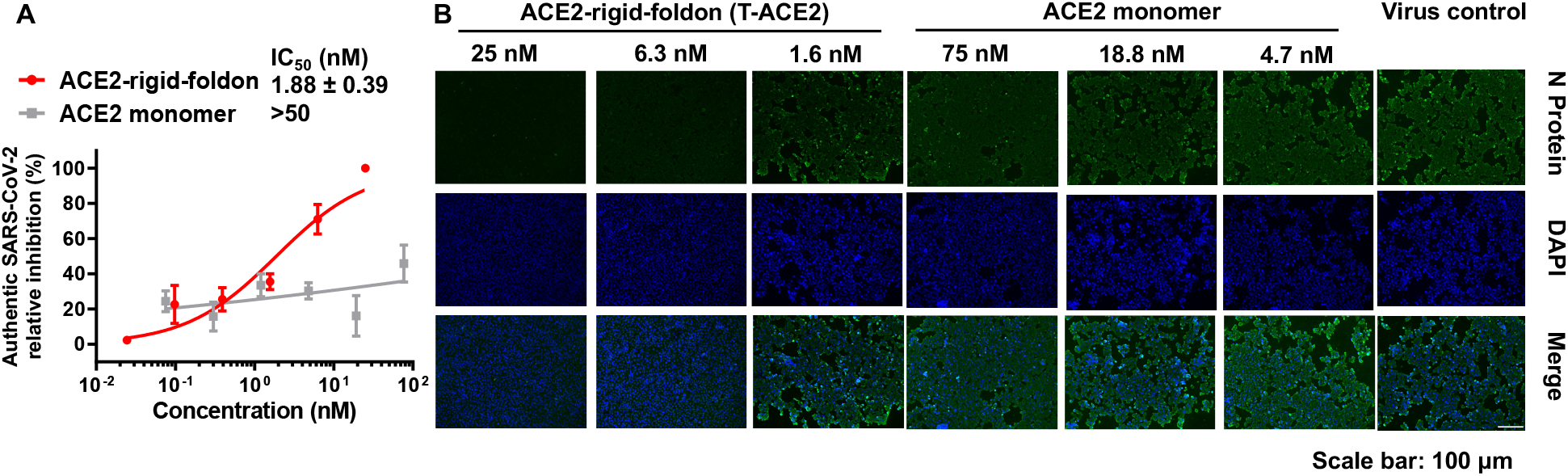
ACE2-rigid-foldon (T-ACE2) inhibition of authentic SARS-CoV-2 virus (n=3). **(A)** Vero E6 cells were infected with authentic SARS-CoV-2, and inhibitory effects were evaluated using quantitative real-time (qPCR). **(B)** Immunofluorescence microscopy of virus infection upon treatment of ACE2-rigid-foldon (T-ACE2) or ACE2 monomer.

We hypothesized that properly designed trimeric ACE2 might engage more than one RBDs from the trimeric spike protein and thus dramatically increase binding affinity through avidity effect. To further test this hypothesis, we cleaved off T-ACE2 C-terminal tag using HRV3C protease, incubated with S-ECD and determined the complex structure using cryo-electron microscopy (cryo-EM) (fig. S6-S8). For simplicity, we still kept the same name T-ACE2 for the C-terminal tag cleaved protein.

Strikingly, in the complex, the spike protein adopts only one conformation: the “three-up” RBD conformation. The complex has a nearly perfect three-fold symmetry. Most importantly, all three RBDs bind to three ACE2s simultaneously. (**Fig. 4**). The binding interactions between ACE2 and RBD are essentially the same as previous studies, and the three copies from the complex align quite well (fig. S8-S9) ^42,43^. Although we couldn’t observe the linker and the trimerization motif, we are relatively confident that the three ACE2s binding to the same spike protein are from the same trimer because of the binding affinity data and virus inhibition data. This spike protein conformation is very different from the previously reported prefusion stabilized spike protein structures where only one or no RBD is in the up position. Recent complex structures between SARS-CoV-2 spike protein and ACE2 monomer from preprints indicate that monomer ACE2 binding can induce conformational changes of spike protein and that some spike protein can have two or three RBDs in the up position to bind up to three ACE2s ^44,45^. In our structure, the unique “three-up” RBD conformation in all the spike proteins should, indeed, be induced by our trimeric ACE2.

**Fig. 4.**
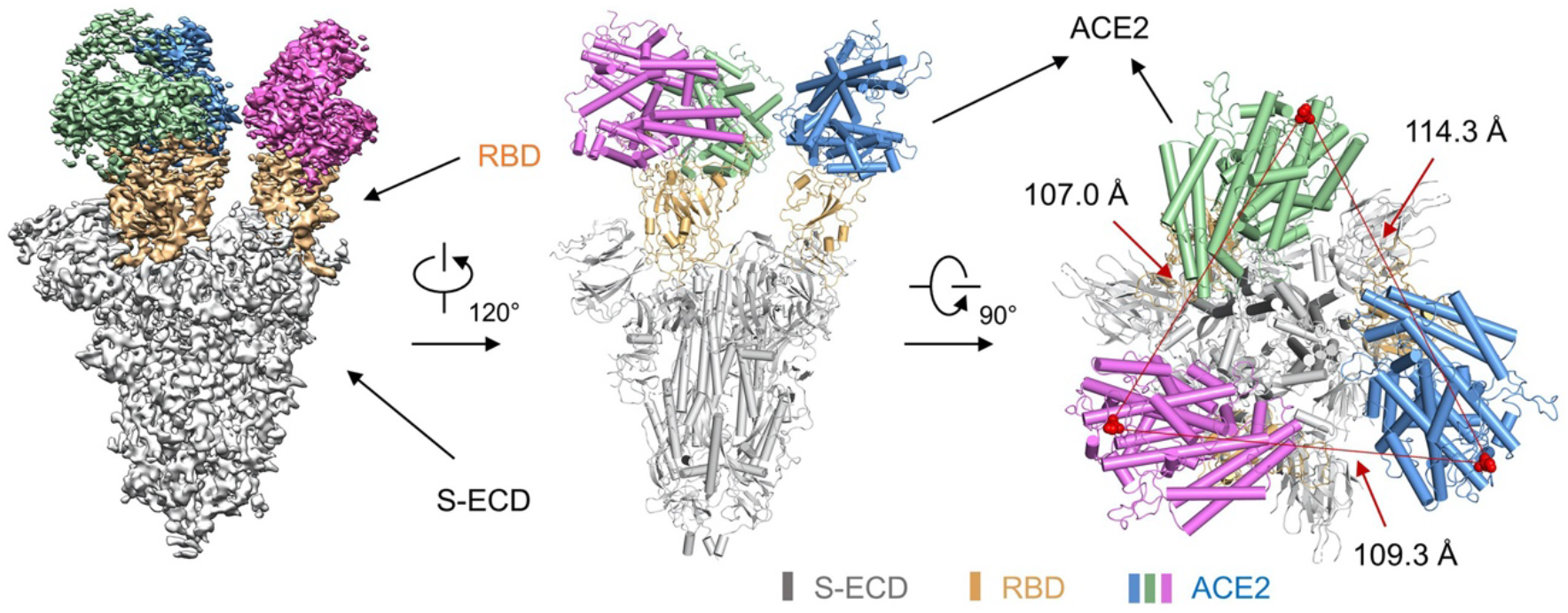
Cryo-EM structure of the T-ACE2/S-ECD complex. The domain-colored cryo-EM map of the complex is shown on the left. Two perpendicular views of the overall structure are shown on the right. The three ACE2 monomers from T-ACE2 are colored blue, green and violet, respectively. The RBDs of the S-ECD are colored orange.

The distance between the C-terminal end of the three ACE2s is around 110 Å (**Fig. 4**). If the trimerization motif sits right in the center, then the ideal linker length between trimerization motif and ACE2 would be around 60 Å, which corresponds to the length of the (GGGGS)3 linker. Thus, the (GGGGS)5 flexible linker in some of our designed proteins is long enough for three ACE2s to bind, but is not optimal. The more rigid (EAAAK)5 linker is shorter than (GGGGS)5 and can effectively separate different functional domains of fusion proteins ^46^. We think the effective length of the (EAAAK)5 linker is probably around 60 Å, making it an optimal linker for T-ACE2. The rigid nature of this (EAAAK)5 linker probably helps to orient ACE2 right around RBD for immediate rebinding even if one ACE2 monomer from T-ACE2 dissociates from the spike protein.

## Discussion

Since the beginning of COVID-19 pandemic, tremendous efforts have been made to develop therapeutics, especially those that utilize neutralizing antibodies. However, the widespread and ongoing crisis of COVID-19 indicates that SARS-CoV-2 will not soon be eliminated, making it prudent to anticipate future mutations notably obviating any current neutralizing antibodies. Moreover, the emergence of COVID-19 after SARS suggests that related coronavirus pandemics might happen in the future. Such events call for therapeutic approaches widely useful for current and future related coronaviruses and mutants.

Several engineered ACE2 proteins bearing different number of mutations and their Ig fusion proteins have been shown to increase spike protein binding affinities and virus inhibitory activities, albeit with reduced or loss of catalytic activities ^28,47^. ACE2-Ig fusion proteins and neutralizing antibodies potentially could have antibody-dependent enhancement (ADE) effect to facilitate virus infection, although such phenomenon still needs further clinical studies ^48,49^. Here, we engineered trimeric ACE2 proteins based on wild-type ACE2 and showed that T-ACE2 could bind spike protein with extremely high affinity to potently inhibit all tested pseudotyped viruses including SARS-CoV-2, SARS-CoV, eight naturally occurring SARS-CoV-2 mutants, two SARSr-CoVs as well as authentic SARS-CoV-2. The rigid linker employed in T-ACE2 was previously injected into mice and didn’t seem to show strong immunogenicity ^50^. The 3HB and foldon trimerization motifs have been observed to cause immunogenicity, but the introduction of glycans could silence the immunogenicity without disrupting the trimer formation ^51^. Carrying these advancements a few steps beyond, the modular design of T-ACE2 demonstrates that other oligomerization motifs and linkers could be further explored to improve properties of T-ACE2 or higher oligomeric ACE2s.

We demonstrated that T-ACE2 could induce the transit of spike protein to a unique “three-up” RBD conformation and bind all three RBDs simultaneously. Whether this T-ACE2-induced spike protein conformation change represents a transition state during virus infection cannot be definitively answered here. Full-length ACE2 protein functions as a dimer ^42^, the two monomers from this ACE2 dimer are related by two-fold symmetry. They are also situated close in space, with the distance between D615 being about 53 Å. Thus, the native dimeric ACE2 is unlikely to engage more than one RBD from the same spike protein without substantial conformational changes. It is however possible that ACE2 dimers on the cell surface might further cluster to induce more RBDs to adopt up conformation and help virus to transit from the prefusion state to the postfusion state.

ACE2 plays an important role in negatively regulating the renin-angiotensin-aldosterone system (RAS) to counterbalance ACE ^52^. Downregulation of ACE2 and elevated plasma angiotensin II level have been observed after SARS-CoV, SARS-CoV-2 or influenza infection, contributing to hyper-activated RAS cascades and ARDS ^18–20,22,52^. Supplementing soluble ACE2 can balance RAS and improve ARDS conditions ^17,21–23,52,53^. These factors all seem to support the beneficial effects of the biological function of ACE2 for treating COVID-19 patients even though further studies are still needed to confer this advantage. Thus, proteins engineered based on wild-type ACE2, such as T-ACE2, can potently and broadly inhibit virus infections, they also have the added benefits of regulating RAS and alleviating ARDS. These potential beneficial effects distinguish proteins like T-ACE2 from neutralizing antibodies. We believe T-ACE2 represents a promising class of proteins to broadly inhibit SARS-CoVs and to treat viruses infected patients. Finally, the extremely high binding affinity between T-ACE2 and spike protein (K_D_ < 1pM) suggests that T-ACE2 could also be useful for virus detection. The fact that T-ACE2 was engineered based on native ACE2 sequence also makes such detection methods widely useful for all SARS-CoVs and related viruses.

## Acknowledgments

We thank the Cryo-EM facility, Supercomputer Center of Westlake University for providing cryo-EM and computation support, respectively. We thank members of the Core Facility of Microbiology and Parasitology (SHMC) and the Biosafety Level 3 Laboratory at Shanghai Medical College of Fudan University, especially Qian Wang, Zhiping Sun, Chengjian Gu, for continuous support. We thank Tian Li for helping protein expression and purifications, we thank Mohamad Sawan, Dan Ma and Huaizong Shen for sharing protein expression, purification and cell culture facilities.

## Author contributions

BD conceived the project. LG prepared all the ACE2 proteins and did binding affinity measurements with help from KZ, MZ and XB. WB and XW did pseudotyped virus inhibition assays under the supervision of BD, LL, YX, and SJ with help from XC and YL. WX, XC and YL did authentic virus inhibition assays under the supervision of YX, DQ and LL. RY, YZ and YL did cryo-EM structure determination under the supervision of QZ. BD, LL, QZ, YX and SJ interpreted the data, wrote and revised the manuscript.

## Data and materials availability

Atomic coordinates and cryo-EM density maps of the S-ECD of SARS-CoV-2 in complex with T-ACE2 (PDB: 7CT5; whole map: EMD-30460, local map of the interface between RBD of SARS-CoV-2 and ACE2: EMD-30461) have been deposited to the Protein Data Bank (http://www.rcsb.org) and the Electron Microscopy Data Bank (https://www.ebi.ac.uk/pdbe/emdb/), respectively.

## Methods

### Protein preparations

To construct trimeric ACE2s, we inserted the linker sequences (GGGGS)5, (EAAAK)5 or GGGS after ACE2(1-615), followed by trimerization motifs, an HRV3C cleavage sequence, an eGFP tag and a His8 tag. Monomeric ACE2 was constructed as ACE2(1-615)-(GGGGS)5-HRV3C-eGFP-His8 for direct comparison.

The ACE2 (accession number: NM_001371415) peptidase domain (1-615) was cloned from the plasmid donated by Peihui Wang’s lab. The genes of 3HB and foldon were synthesized by Genewiz, Suzhou, China. All the gene fragments were assembled by the Gibson assembly kit (Cat.C112-01, Vazyme). The assembled fragments were subcloned in pEGFP-N1 for expression. The cloned plasmids were transformed into E.coli DH5α for amplification. Amplified plasmids were extracted using the GoldHi EndoFree Plasmid Maxi Kit (Cat. CW2104M, CWBio).

HEK 293F cells (Invitrogen) were cultured in Freestyle medium (Gibco, Lot.2164683) at 37 °C under 6% CO2 in a CRYSTAL shaker (140 rpm). The cells were transiently transfected with ACE2 plasmids and polyethylenimine (PEI) (Polysciences, Cat.24765-1) when the cell density reached approximately 1.0×10^6^/mL. 1 mg of plasmid was premixed with 2.6 mg PEI in 50 mL of fresh medium for 15 minutes before adding to one liter of cell culture. The transfected cells were cultured for 96 hours before harvesting.

For purification of ACE2 proteins, the cell supernatants were harvested by centrifugation at 1000×g for 5 minutes. Then the supernatants were loaded on Ni-NTA beads (Smart-Lifesciences, Cat. SA004100) and washed with washing buffer (5 mM imidazole, 1 × PBS). Proteins were then eluted with elution buffer (50 mM imidazole, 1 × PBS).

The eluted proteins were concentrated and subjected to size-exclusion chromatography (Superose 6 Increase 10/300 GL, GE Healthcare) in the PBS buffer. The peak fractions were collected and concentrated. The proteins were then analyzed by size exclusion chromatography (AdvanceBio SEC 300Å) in PBS buffer pH 7.4. The standard proteins were purchased from GE (fig S2).

To remove C-terminal tags of ACE2 proteins, 16 μg HRV3C protease (expressed and purified in house) was add to 1mg ACE2 protein and incubated at 4 °C overnight, followed by size-exclusion chromatography (Superose 6 Increase 10/300 GL, GE Healthcare) purification and analysis.

### Binding affinity measurement using ELISA assays

#### Determination of optimal S-ECD loading

96-well ELISA plates (JET BIOFIL, #FEP-100-096) were coated with 50 μL per well of different S-ECD protein concentrations (fig S3) in coating buffer (NCM Biotech, #E30500) overnight at 4 °C. Plates were washed with phosphate-buffered saline with 0.1% Tween-20 (PBST) four times and then blocked with 2% bovine serum albumin (BSA, Sigma, #B2064-50G) in PBST for 2 hours at room temperature. After blocking, the plates were washed with PBST four times and then incubated with 70 μL per well of ACE2 monomer in PBST for 2 hours at 37°C. Plates were washed with PBST four times then incubated with 70 μL per well of 1:2,000 dilution of Anti-GFP antibody (Rabbit PAb, Sino Biological, #13105-RP01) for 1 h at 37 °C. Plates were again washed four times, followed by incubation with 70 μL per well of 1:10,000 dilution of HRP-conjugated Goat Anti-Rabbit IgG (Beyotime, #A0208) for 1 hour at 37 °C. After final washing, 100 μL per well of TMB single-component substrate solution were added to the plates (Solarbio, #PR1200), and the reaction was stopped by the addition of 50 μL per well of 1M hydrochloric acid. The absorbance at 450 nm was measured on a Microplate reader (Thermo, Varioskan LUX). From this experiment, we decided to load 3 μg/mL S-ECD for ACE2 protein binding measurement.

#### Binding measurements

To determine the binding affinities of different ACE2 proteins, 96-well ELISA plates (JET BIOFIL, #FEP-100-096) were coated with 50 μL per well of S-ECD (3 μg/mL) in coating buffer (NCM Biotech, #E30500) overnight at 4 °C. Plates were washed with phosphate-buffered saline with 0.1% Tween-20 (PBST) four times and then blocked with 2% bovine serum albumin (BSA, Sigma, #B2064-50G) in PBST for 2 hours at room temperature. After blocking, the plates were washed with PBST four times and then incubated with 70 μL per well of series diluted ACE2 samples in PBST for 2 hours at 37°C. Plates were washed with PBST four times and then incubated with 70 μL per well of 1:2,000 dilution of Anti-GFP antibody (Rabbit PAb, Sino Biological, #13105-RP01) for 1 hour at 37 °C. Plates were again washed four times, followed by incubation with 70 μL per well of 1:10,000 dilution of HRP-conjugated Goat Anti-Rabbit IgG (Beyotime, #A0208) for 1 hour at 37 °C. After final washing, 100 μL per well of TMB single-component substrate solution were added to the plates (Solarbio, #PR1200), and the reaction was stopped by the addition of 50 μL per well of 1M hydrochloric acid. The absorbance at 450 nm was measured on a microplate reader (Thermo, Varioskan LUX).

### Binding affinity determination using bio-layer interferometry (BLI)

#### Protein biotinylation

Purified S-ECD protein was biotinylated at a theoretical 1:3 molar ratio with EZ-Link NHS-PEG12-Biotin (Thermo Fisher Scientific, CAT#: 21313) according to the manufacturer’s instructions. The unreacted biotin was removed by ultrafiltration with an Amicon column (30 KDa MWCO, Millipore, CAT: UFC5010BK).

#### Kinetics analyses

For kinetics analyses, S-ECD was captured on streptavidin biosensors. Biotinylated S-ECD was diluted to 20 μg/mL in dilution buffer (PBS with 0.02% Tween 20 and 0.1% BSA). Then sensor baselines were equilibrated in the dilution buffer for 90 seconds. Next, the S-ECD was loaded until the thickness signal was 0.6 nm or 0.3 nm (low loading). After loading, the sensors were washed for 60 seconds in the dilution buffer. The sensors were then immersed into wells containing ACE2 proteins for 100 seconds (association phase), followed by immersion in dilution buffers for an additional 300 seconds (dissociation phase). The background signal was measured using a reference sensor with S-ECD loading but no ACE2 protein binding and was subtracted from the corresponding ACE2 binding sensor. Curve fitting was performed using a 1:1 binding model and the ForteBio data analysis software. Mean kon, koff, K_D_ values were determined by averaging all binding curves that matched the theoretical fit with an R2 value of 0.95.

### Cell lines, plasmids construction and virus

Human hepatoma Huh-7 cells were purchased from the Cell Bank of the Chinese Academy of Science (Shanghai, China). Human primary embryonic kidney cells (293T) (CRL-3216™) and African green monkey kidney Vero-E6 (CRL-1586™) were obtained from the American Type Culture Collection (ATCC). These cells were cultured with Dulbecco’s Modified Eagle’s Medium (DMEM) containing 10% Fetal bovine serum (FBS), 100 mg/mL streptomycin, and 100 U/mL penicillin at 37 °C under 5% CO_2_.

The envelope-encoding plasmids of SARS-CoV-2-S, SARS-CoV-S, and SARSr-CoV-S (Rs3367 and WIV1) and luciferase-expressing vector (pNL4-3.Luc.R-E-) were maintained in house. The plasmids encoding mutant SARS-CoV-2-S (V341I, F342L, V367F, R408I, A435S, G476S, V483A and D614G) were constructed using a site mutation kit (Yeasen, China) and confirmed by sequencing.

SARS-CoV-2 (SARS-CoV-2 / SH01 / human / 2020 / CHN, GenBank No. MT121215) was isolated from a COVID-19 patient in Shanghai, China. The virus was purified and propagated in Vero-E6 cells, then stocked at −80 °C. Viral titer was measured by the 50% Tissue culture infective dose (TCID50) method. All experiments involving live SARS-CoV-2 virus were performed in Biosafety Level 3 Laboratory (BSL-3), Fudan University.

### Pseudotyped virus inhibition

#### Packaging pseudotyped SARS-CoV-2, mutant SARS-CoV-2, SARS-CoV, and SARSr-CoVs

These pseudoviruses were generated according to previous studies ^54,55^. Briefly, the envelope-encoding plasmid (20 μg) and pNL4-3.Luc.R-E-(10 μg) were cotransfected into 293T cells cultured in a 10 cm cell culture dish using Vigofect transfection reagent (Vigorous Biotechnology, China). After 10 hours, the cell culture medium was changed with fresh DMEM containing 10% FBS. Supernatants containing pseudovirus were harvested 48 hours later, filtered with a 0.45 μm filter (Millipore), and used for single-cycle infection.

#### Inhibition of Pseudotyped SARS-CoV-2, SARS-CoV-2 mutants, SARS-CoV, and SARSr-CoVs infections

The pseudotyped viruses inhibition assays were conducted as previously described ^54,55^. Briefly, Huh-7 cells were seeded into the 96-well cell culture plate at 1×10^4^ per well and cultured for 12 hours. The recombinant proteins were diluted with FBS-free DMEM and mixed with pseudotyped viruses, incubated at 37 °C for 30 minutes, and added to Huh-7 cells. After 12 hours of infection, the culture medium was replaced with fresh DMEM containing 10% FBS, and cells were cultured for an additional 48 hours. Then cells were lysed with Cell Lysis Buffer (Promega, Madison, WI, USA), and the luciferase activity was detected using the Luciferase Assay System (Promega, Madison, WI, USA), all data were analyzed using Prism Graphpad.

### Authentic SARS-CoV-2 virus inhibition or authentic SARS-CoV-2 neutralization

The live SARS-CoV-2 inhibition assay was performed as previously described ^56^. Briefly, Vero-E6 cells were seeded into the 96-well cell culture plate at 3×10^4^ per well and cultured for 12 hours. Recombinant proteins were diluted with FBS-free DMEM, mixed with 100 TCID50 of SARS-CoV-2, and incubated at 37°C for 1 hour. Then, the protein-virus mixtures were added to Vero-E6 cells and incubated at 37°C for 1 hour. After removing the mixtures, cells were cultured with fresh DMEM containing 2% FBS for another 48 hours. Then, the supernatants were collected to detect viral RNA titer. The cells were fixed to perform immunofluorescence analysis. After fixing with 4% paraformaldehyde, the cells were permeabilized by 0.2% Triton X-100 and blocked with 3% BSA for 1 hour. Then the SARS-CoV-2 Nucleocapsid Antibody (1:500) (Sino Biological) was added to cells and reacted at 4°C overnight. Finally, the cells were incubated with Alexa Fluor 488 Goat anti-Rabbit IgG (1:500) (Invitrogen, USA) at 37°C for 1 hour. The nuclei were stained with NucBlue™ Live ReadyProbes™ Reagent (Thermo Fisher Scientific, USA) and imaged with fluorescence microscopy.

#### RNA extraction and Quantitative Real-time PCR (qPCR) assay

Total viral RNA in supernatants were extracted using Trizol LS reagent (Invitrogen, USA), according to the manufacturer’s manual. Then qPCR was conducted with a One-Step PrimeScrip RT-PCR Kit (Takara, Japan), following the manufacturer’s instructions. qPCR reaction was performed with the program of 95 °C for 10 seconds, 42°C for 5 minutes; 40 cycles of 95 °C for 5 seconds, 50 °C for 30 seconds, 72 °C for 30 seconds on Bio-Rad CFX96. Viral loads were determined by a standard curve prepared with a plasmid containing SARS-CoV-2 nucleocapsid protein (N) gene (purchased form BGI, China). Primers and probe targeting SARS-CoV-2 N gene were ordered from Genewiz (Suzhou, China), and the sequences were as follows: SARS-CoV-2-N-F: GGGGAACTTCTCCTGCTAGAAT, SARS-CoV-2-N-R: CAGACATTTTGCTCTCAAGCTG, SARS-CoV-2-N-probe: 5’-FAM-TTGCTGCTGCTTGACAGATT-TAMRA-3’.

### Cryo-EM sample preparation

Purification of the extracellular domain (ECD) (Genebank ID: QHD43416.1) (1-1208 a.a) of S protein was as previously reported ^57^. For structure determination, we cleaved the C-terminal tag of T-ACE2 using HRV3C protease. Purified S-ECD was mixed with the T-ACE2 at a molar ratio of about 1:2 for one hour at 4 °C. The mixture was subjected to size-exclusion chromatography (Superose 6 Increase 10/300 GL, GE Healthcare) in buffer containing 25 mM Tris (pH 8.0), 150 mM NaCl. Peak fractions of S-ECD in complex with T-ACE2 were collected for EM analysis.

The peak fractions of the complex were concentrated to about 1.5 mg/mL and mixed with 0.05% Octyl Maltoside, Fluorinated (Anatrace) before application to the grids. Aliquots (3.3 μL) of the protein complex were placed on glow-discharged holey carbon grids (Quantifoil Au R1.2/1.3). The grids were blotted for 2.5 s or 3.0 s and flash-frozen in liquid ethane cooled by liquid nitrogen with Vitrobot (Mark IV, Thermo Scientific). The cryo-EM samples were transferred to a Titan Krios operating at 300 kV equipped with Cs corrector, Gatan K3 Summit detector and GIF Quantum energy filter. Movie stacks were automatically collected using AutoEMation ^58^ with a slit width of 20 eV on the energy filter and a defocus range from −1.2 μm to −2.2 μm in super-resolution mode at a nominal magnification of 81,000×. Each stack was exposed for 2.56 s with an exposure time of 0.08 s per frame, resulting in a total of 32 frames per stack. The total dose rate was approximately 50 e^-^/Å^2^ for each stack. The stacks were motion corrected with MotionCor2 ^59^ and binned 2-fold, resulting in a pixel size of 1.087 Å/pixel. Meanwhile, dose weighting was performed ^60^. The defocus values were estimated with Gctf ^61^.

### Data processing

Particles were automatically picked using Relion 3.0.6 ^62–65^ from manually selected micrographs. After 2D classification with Relion, good particles were selected and subjected to two cycles of heterogeneous refinement without symmetry using cryoSPARC ^66^. The good particles were selected and subjected to Non-uniform Refinement (beta) with C1 symmetry, resulting in the 3D reconstruction for the whole structures that were further subjected to 3D classification, 3D auto-refinement and post-processing with Relion. For interface between RBD and ACE2, the datasets were subjected to focused refinement with adapted mask on each RBD and ACE2 subcomplex to improve the map quality. Then the dataset of three RBD and ACE2 sub-complexes were combined and subjected to focused refinement with Relion, resulting in the 3D reconstruction of better quality on the interface between S-ECD and ACE2.

The resolution was estimated with the gold-standard Fourier shell correlation 0.143 criterion ^67^ with high-resolution noise substitution ^68^. Refer to (fig S6-8) and Supplemental Table S1 for details of data collection and processing.

### Model building and structure refinement

For model building of the complex of S-ECD with ACE2, the atomic model of the published structure S-ECD (PDB ID: 7C2L) and ACE2 molecular (PDB ID: 6M18) were used as templates, which were molecular dynamics flexible fitted (MDFF) ^69^ into the whole cryo-EM map of the complex and the focused-refined cryo-EM map of the RBD-ACE2 subcomplex, respectively. The fitted atomic models were further manually adjusted with Coot ^70^. Each residue was manually checked with the chemical properties taken into consideration during model building. Several segments, the corresponding densities of which were invisible, were not modeled. Structural refinement was performed in Phenix ^71^ with secondary structure and geometry restraints to prevent overfitting. To monitor the potential overfitting, the model was refined against one of the two independent half maps from the gold-standard 3D refinement approach. Then, the refined model was tested against the other map. Statistics associated with data collection, 3D reconstruction and model building were summarized in Table S1.

---

**Fig. S1.**
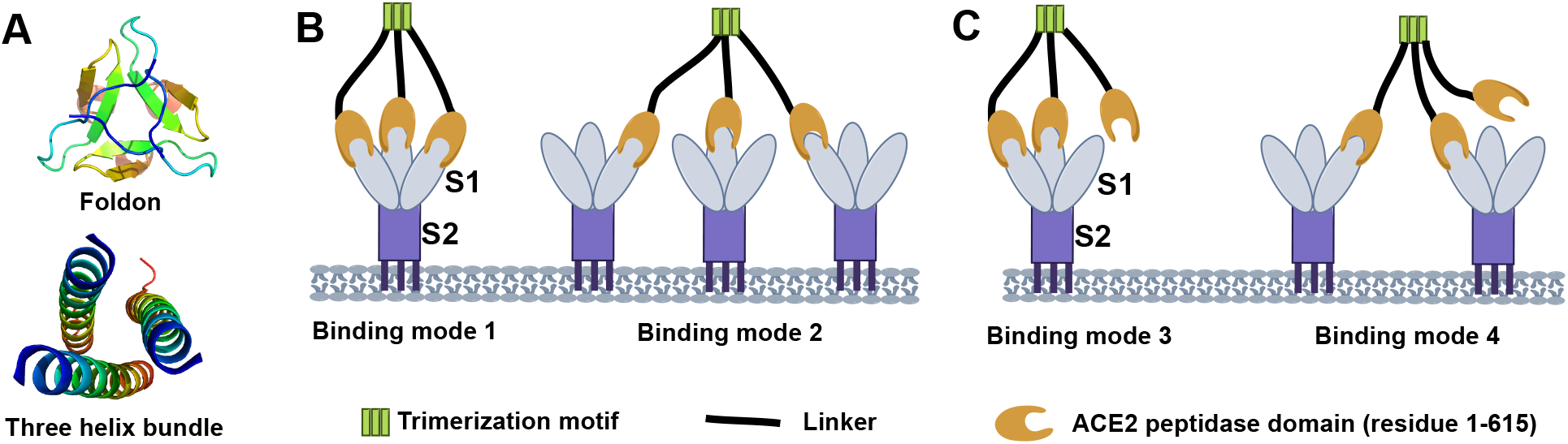
ACE2 trimerization strategy and potential interactions between trimeric ACE2 and spike protein trimer. **(A)** Structures of the two ACE2 trimerization motifs. **(B)** Each ACE2 trimer can engage three RBDs either from the same spike protein (mode 1) or different spike proteins (mode 2). **(C)** Only two ACE2s from the trimer can engage two RBDs either from the same spike protein (mode 3) or different spike proteins (mode 4).

**Fig. S2.**
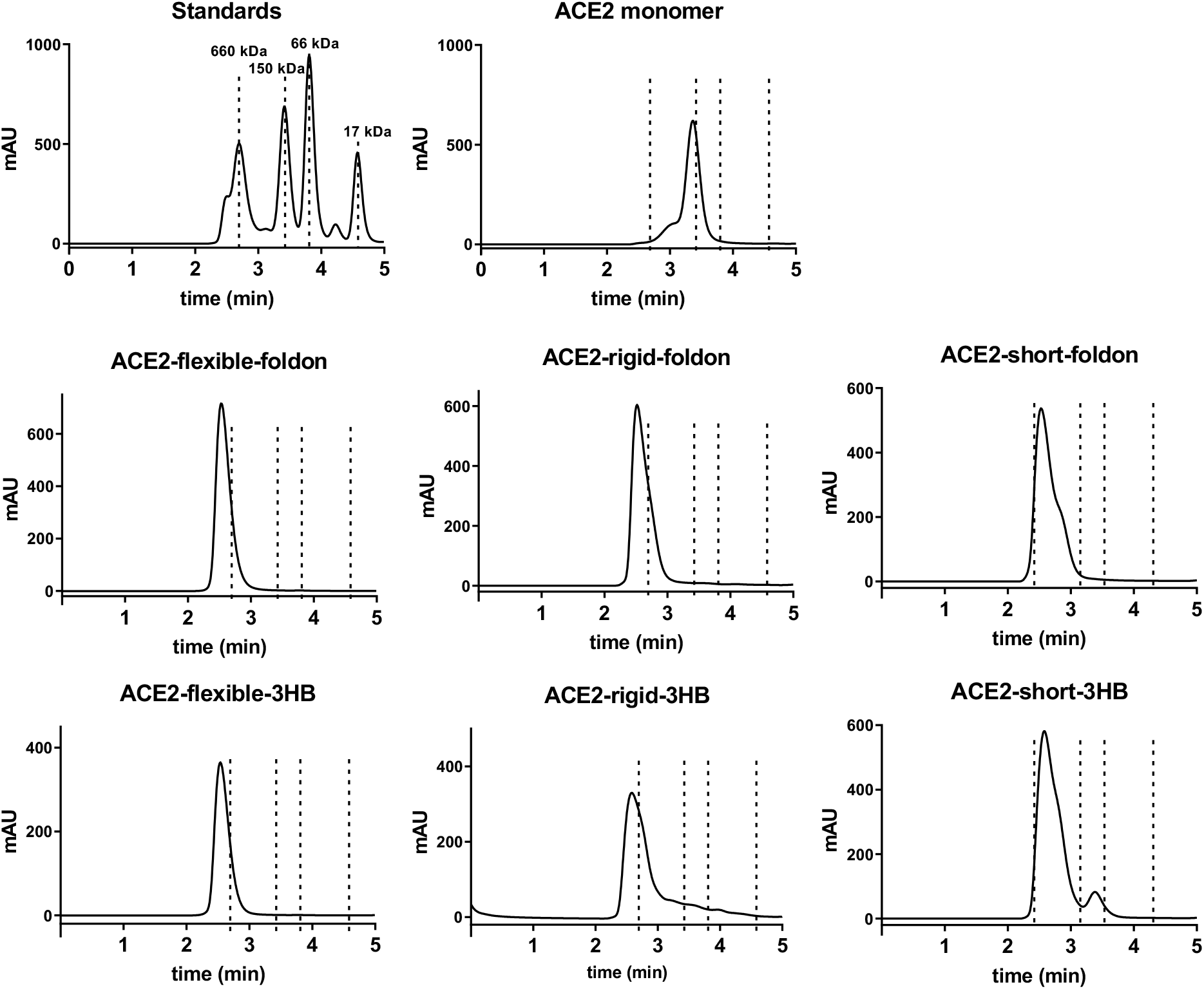
Size-exclusion chromatography analyses of ACE2 proteins.

**Fig. S3.**
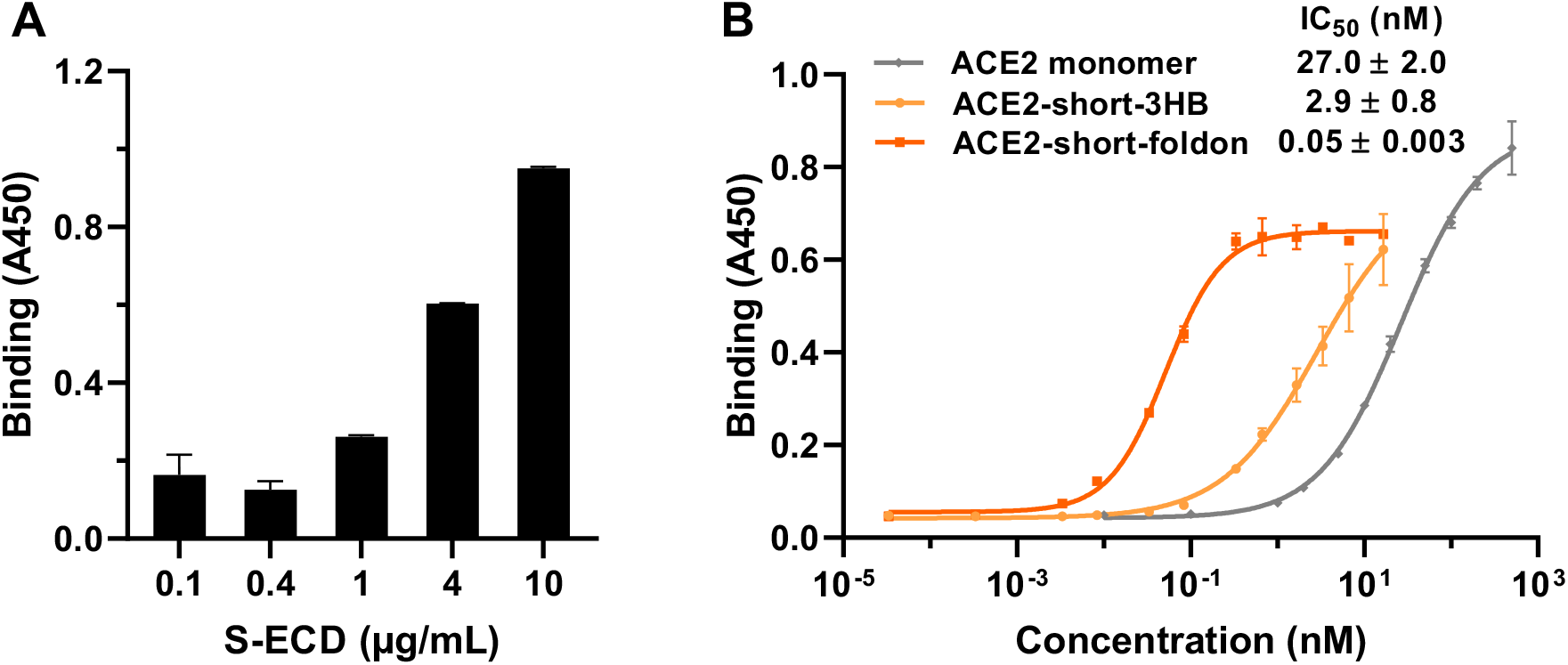
ELISA binding measurements. **(A)** S-ECD loading amount optimization. **(B)** Short linker ACE2 proteins binding affinities to S-ECD determined in ELISA assay.

**Fig. S4.**
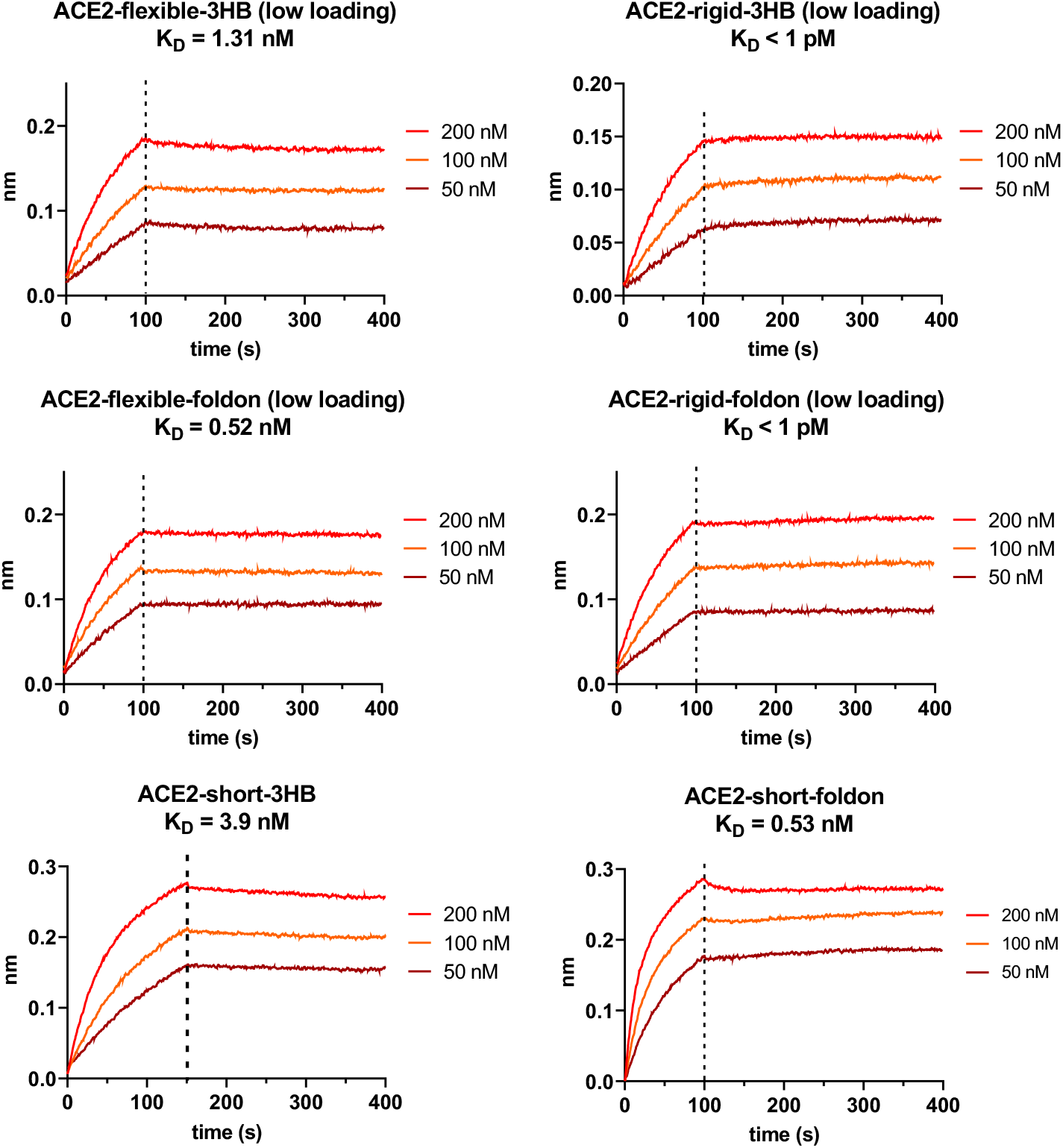
Binding affinities measurement between ACE2 proteins and SARS-CoV-2 spike protein ectodomain (S-ECD). Low loading means S-ECD was loaded at thickness signal of 0.3 nm, whereas normal loading is thickness signal of 0.6 nm.

**Fig. S5.**
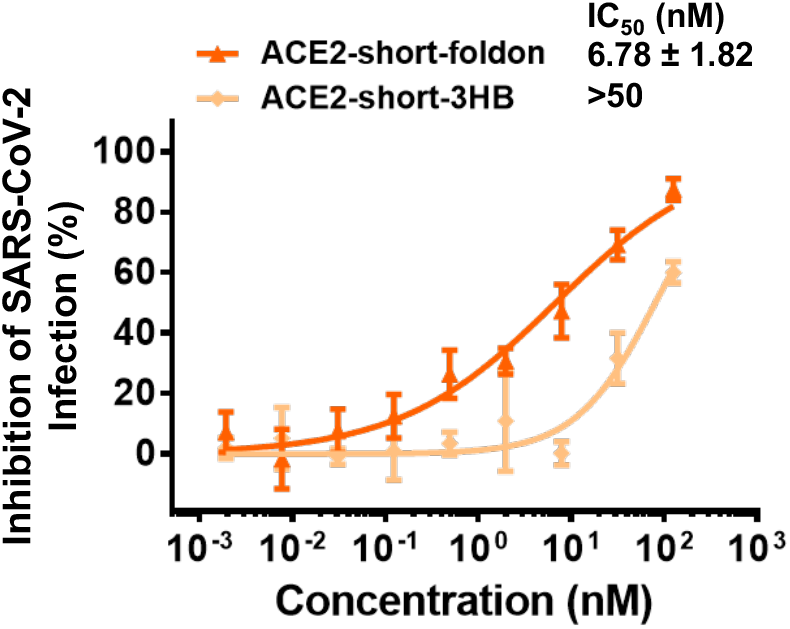
Short-linker ACE2 proteins inhibition of SARS-CoV-2 pseudotyped virus.

**Fig. S6.**
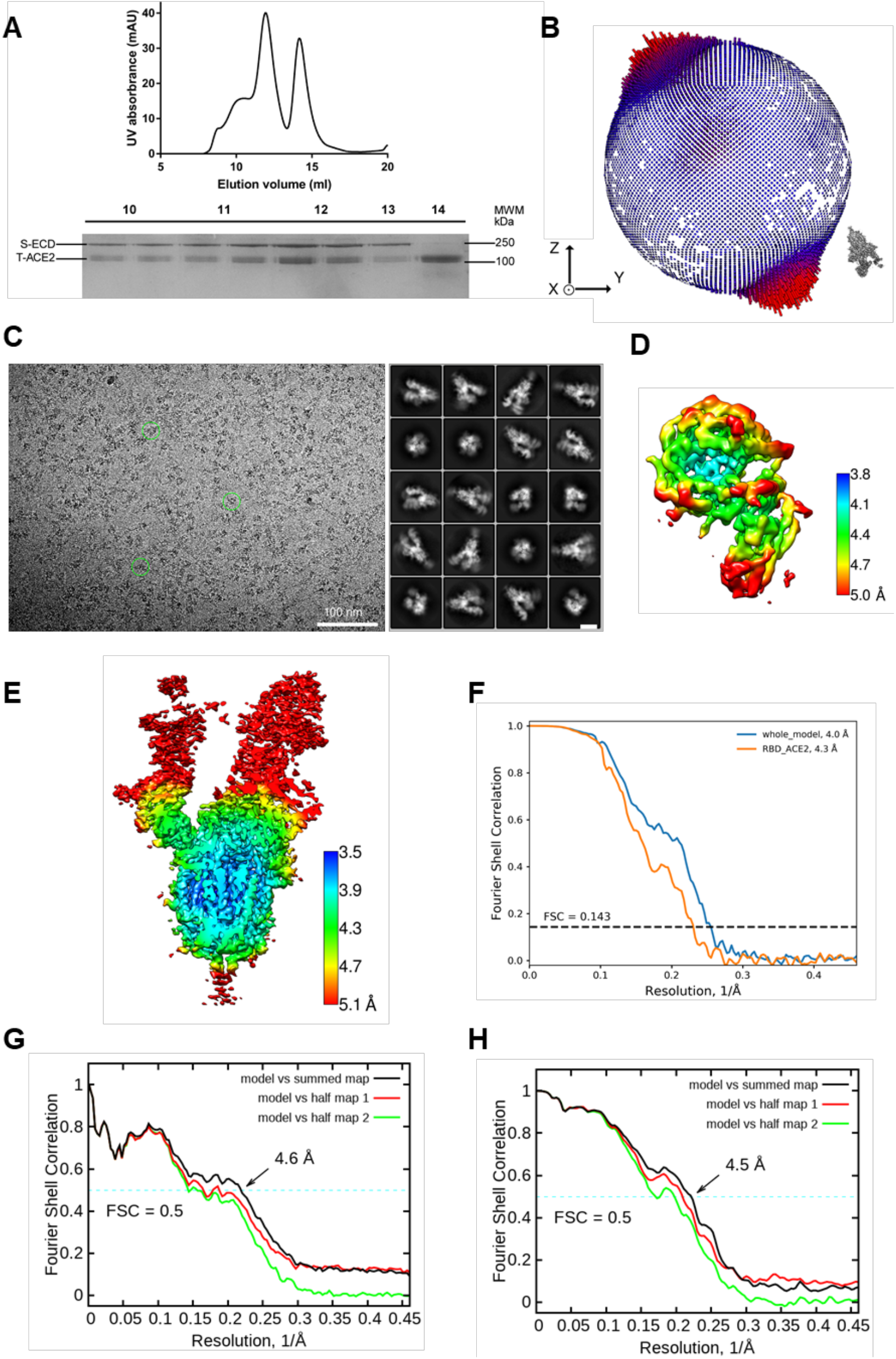
Cryo-EM analysis of S-ECD in complex with ACE2. **(A)** Representative SEC purification profile of the S-ECD in complex with T-ACE2. **(B)** Euler angle distribution in the final 3D reconstruction of S-ECD in the SARS-CoV-2/T-ACE2 complex. **(C)** Representative cryo-EM micrograph and 2D class averages of cryo-EM particle images. The scale bar in 2D class averages is 10 nm. **(D)** and **(E)** Local resolution maps for the 3D reconstruction of the RBD-ACE2 subcomplex and overall structure, respectively. **(F)** FSC curve of the overall structure (blue) and RBD-ACE2 subcomplex (orange). **(G)** FSC curve of the refined model of S-ECD of SARS-CoV-2 bound with ACE2 complex versus the overall structure against which it is refined (black), the refined model against the first half of the map versus the same map (red); and the refined model against the first half of the map versus the second half map (green). The small difference between the red and green curves indicates that the refinement of the atomic coordinates did not allow enough for overfitting. **(H)** FSC curve of the refined model of RBD-ACE2 subcomplex is the same as **(G)**.

**Fig. S7.**
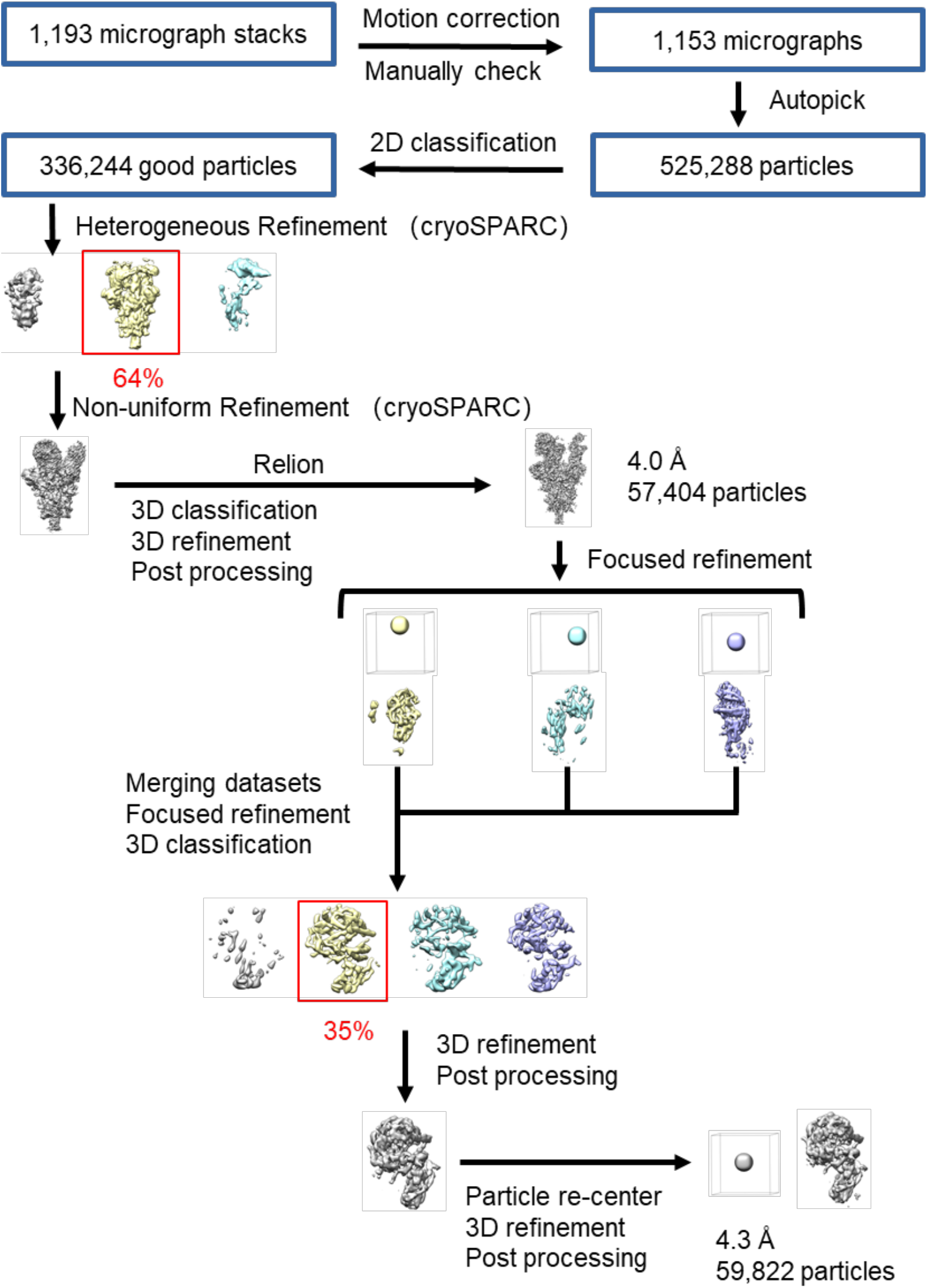
Flowchart for cryo-EM data processing. Please refer to the ‘Data Processing’ section in Methods for details.

**Fig. S8.**
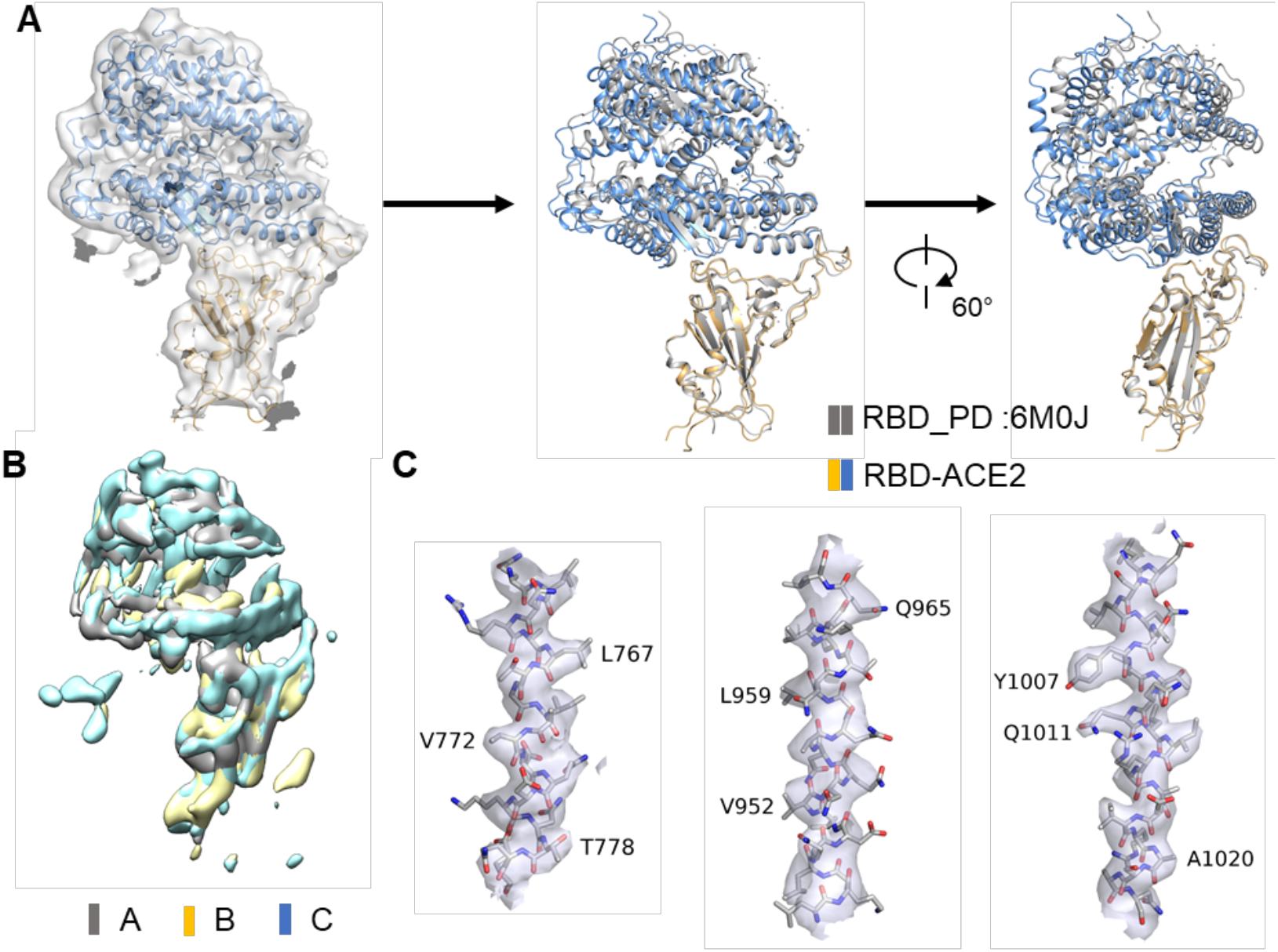
Structural analysis and representative cryo-EM map densities of S-ECD in complex T-ACE2. **(A)** Structural alignment in the interface of RBD and ACE2 with the RBD-PD complex previously reported (PDB ID: 6M0J) with a root mean squared deviation of 0.776 Å over 178 pairs of Cα atoms. **(B)** Superposition in local map of RBD-ACE2 subcomplex for three protomers, indicating no difference among three maps. **(C)** Representative cryo-EM map densities of S-ECD in complex T-ACE2. All densities are shown at the threshold of 5 σ.

**Fig. S9.**
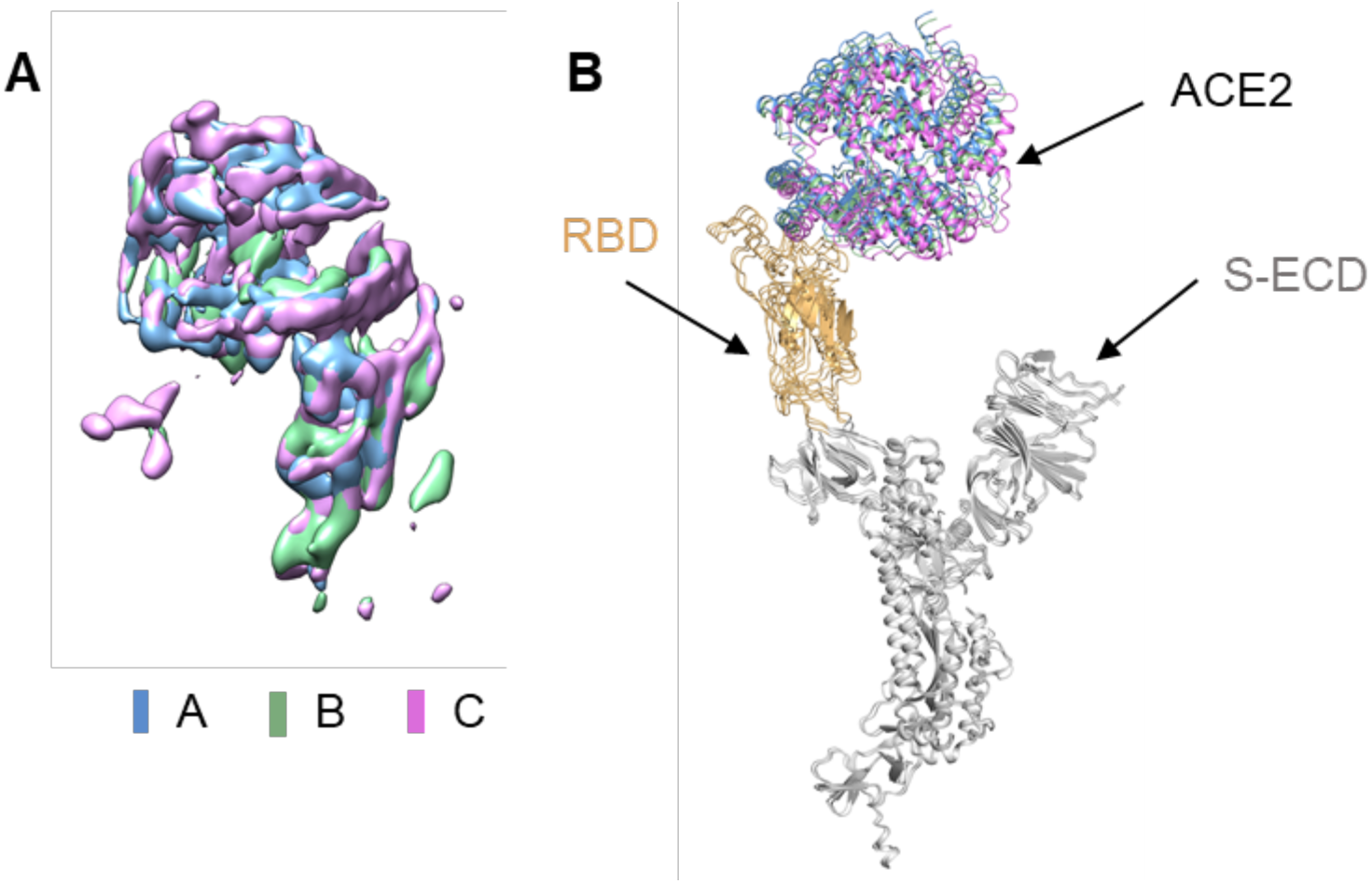
Structural alignment of three protomer for S-ECD in complex ACE2. **(A)** Superposition in local map of RBD-ACE2 sub-complex for three protomer, which has no difference among three maps. The three ACE2 are colored blue, green and violet, respectively. **(B)** Structural alignment of three monomer of S-ECD in complex ACE2.

**Table S1.**
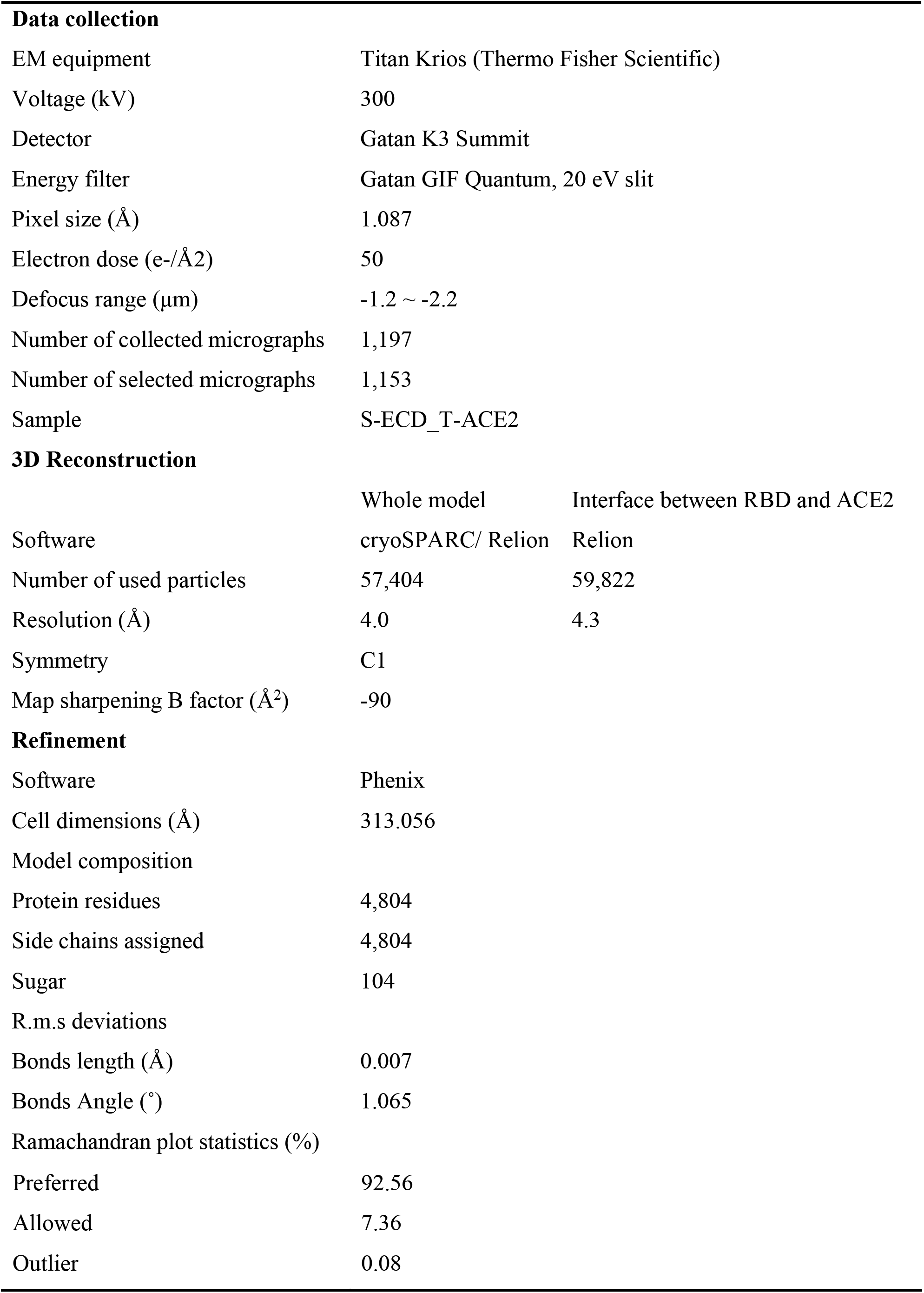
Cryo-EM data collection and refinement statistics.

|  |  |  |
| --- | --- | --- |
| **Data collection** |  |  |
| EM equipment | Titan Krios (Thermo Fisher Scientific) |  |
| Voltage (kV) | 300 |  |
| Detector | Gatan K3 Summit |  |
| Energy filter | Gatan GIF Quantum, 20 eV slit |  |
| Pixel size (Å) | 1.087 |  |
| Electron dose (e-/Å2) | 50 |  |
| Defocus range (µm) | -1.2 ~ -2.2 |  |
| Number of collected micrographs | 1,197 |  |
| Number of selected micrographs | 1,153 |  |
| Sample | S-ECD_T-ACE2 |  |
| **3D Reconstruction** |  |  |
|  | Whole model | Interface between RBD and ACE2 |
| Software | cryoSPARC/ Relion | Relion |
| Number of used particles | 57,404 | 59,822 |
| Resolution (Å) | 4.0 | 4.3 |
| Symmetry | C1 |  |
| Map sharpening B factor (Å²) | -90 |  |
| **Refinement** |  |  |
| Software | Phenix |  |
| Cell dimensions (Å) | 313.056 |  |
| Model composition |  |  |
| Protein residues | 4,804 |  |
| Side chains assigned | 4,804 |  |
| Sugar | 104 |  |
| R.m.s deviations |  |  |
| Bonds length (Å) | 0.007 |  |
| Bonds Angle (°) | 1.065 |  |
| Ramachandran plot statistics (%) |  |  |

|  |  |
| --- | --- |
| Preferred | 92.56 |
| Allowed | 7.36 |
| Outlier | 0.08 |

